# Dynamic arm movements attenuate perceptual distortion of visual vertical induced during prolonged whole-body tilt

**DOI:** 10.1101/2020.03.27.011593

**Authors:** Keisuke Tani, Shinji Yamamoto, Yasushi Kodaka, Keisuke Kushiro

**Affiliations:** Laboratory of Psychology, Hamamatsu University School of Medicine, 1-20-1 Handayama, Higashi-ku, Hamamatsu, Shizuoka 431-3192, Japan; Faculty of Sport Sciences, Nihon Fukushi University, Okuda, Mihama-cho, Chita-gun, Aichi 470-3295, Japan; Automotive Human Factors Research Center, National Institute of Advanced Industrial Science and Technology, 1-1-1 Umezono, Tsukuba, Ibaraki 305-8560, Japan; Graduate School of Human and Environmental Studies, Kyoto University, Yoshidanihonmatsu-cho, Sakyou-ku, Kyoto 606-8501, Japan

**Keywords:** Spatial orientation, Arm movement, Gravity, Subjective visual vertical (SVV), Subjective postural vertical (SPV), Prolonged whole-body tilt

## Abstract

Additional gravitational cues generated by active body movements may play a role in the perception of gravitational space, but no experimental evidence has been shown on this. To investigate this possibility, we evaluated how arm movements made against gravity influenced the perceptual distortion of visual and postural vertical induced by prolonged whole-body tilt. In Experiment 1, participants were asked to perform static or dynamic arm movements during prolonged whole-body tilt and we assessed their effects on subjective visual vertical (SVV) at the tilt position (*during-tilt* session) and after tilting back to the upright position (*post-tilt* session). In Experiment 2, we evaluated how static or dynamic arm movements during prolonged tilt subsequently affected the subjective postural vertical (SPV). In Experiment 1, we observed that prolonged tilt induced the SVV shifts toward the side of body tilts in both sessions. The prolonged tilt-induced SVV shifts effectively decreased when performing dynamic arm movements in the *during-tilt* session, but not in the *post-tilt* session. In Experiment 2, the SPV shifted toward the side of prolonged body tilt, which was not significantly influenced by the performance of static or dynamic arm movements. Results of the *during-tilt* session suggest that the central nervous system utilizes additional cues generated by dynamic body movements for the perception of the visual vertical.

## Introduction

To appropriately achieve aimed actions in the gravitational field, the central nervous system (CNS) plans and executes movements based on spatial information, such as the direction of gravity and body. Therefore, accurately perceiving these directions is an important factor determining the quality of motor performance. Previous studies support this assumption by demonstrating that stroke patients, having impaired perception of their directions, experience difficulty in postural control of sitting and standing [1–6].

To build the internal representation of the gravitational direction, the CNS integrates multisensory signals from visual, somatosensory, and vestibular organs with weighting based on reliability [7–9]. One common method to evaluate the perception of the gravitational direction is the subjective visual vertical (SVV) adjustment, where participants are asked to adjust a visual line to the perceived vertical [10]. Although the SVV closely coincides with the actual gravitational vertical in the upright position, the estimation error in other positions depends on the body tilt angle [11–14]. Specifically, for a relatively small tilt angle (< 60°), SVV typically shifts toward the opposite direction of body tilt [15], whereas it shifts toward the direction of body tilt for a larger tilt angle (> 60°) [16]. Another method of assessing the perception of the gravitational direction is the subjective postural vertical (SPV) task, in which participants are asked to indicate their body’s vertical position while being inclined from one tilted side to the other [7]. It is known that SPV is affected by the direction and angle of the initial body tilt [7].

The perception of gravitational direction is influenced by maintaining the body in an inclined posture for a certain time, referred to as a prolonged tilt. The SVV gradually shifts toward the body tilt side during a prolonged tilt [17–22] and remains deviated toward the previously tilted side even after a return to the upright position (i.e. after-effect) [18,20,21]. Likewise, after a prolonged tilt, the SPV shifts toward the prolonged tilt side [23–26]. These time-dependent changes in the SVV and SPV could be mainly attributed to sensory adaptation. For instance, Fernandez and Goldberg [27] showed that the otolith afferent firing rate in primates gradually decreased in the roll head-tilt position. Other studies suggest the possibility that somatosensory adaptation derived from trunk receptors may also contribute to the SVV shifts during prolonged tilt [11, 22]. The angles of the head and body relative to gravity would be sensed to be smaller due to vestibular and somatosensory adaptation, leading to shifts of the perceived direction of gravity toward the prolonged tilt side [28].

The purpose of this study was to investigate how active arm movements during prolonged tilt influence the perception of the gravitational direction. Performing arm movements against gravity generates additional information, such as proprioceptive feedback from muscle spindles, skin and joint receptors, and the Golgi tendon organ, and efferent copy [29], which would provide cues about the gravitational force on the arm. Moreover, the gravitational torque on the shoulder of an extended arm during arm lifting depends on the arm’s position relative to gravity [30]. Based on these findings, we anticipated that the gravitational cues generated by arm movements would play a role in estimating the gravitational direction. If the CNS utilizes the additional cues to internally estimate gravitational space, the decrease of sensed head/body tilt angle due to sensory adaptation may be compensated by the execution of arm movements during prolonged tilt, resulting in suppressing the alteration of the perceived direction of gravity. Contrary to this hypothesis, several studies have demonstrated that the estimation of body tilt orientation [31] or earth-horizontal direction [32] is not influenced by the execution of arm movements. In these studies, however, participants performed the estimation task immediately after the body was tilted or while the body was being tilted. Thus, sensory adaptation would not have occurred, making it difficult to observe the effects of arm movements on the perception of gravitational space. To address this possibility, the present study evaluated whether or how static or dynamic arm movements during prolonged tilt influenced the perception of visual vertical (Experiment 1) and postural vertical (Experiment 2).

## Materials and Methods

### Experiment 1

#### Participants

Fifteen right-handed healthy volunteers (13 males and 2 females, aged 19-33 years) participated in this experiment after giving written informed consent. All of them had normal vision and no neurological, muscular, or cognitive disorders. This study was approved by the Ethics Committee of the Graduate School of Human and Environmental Studies, Kyoto University, and was conducted in accordance with the Declaration of Helsinki (2013).

#### Apparatus

The participants sat on a seat (RSR-7 KK100, RECARO Japan, Japan) mounted on a tilt-table in a completely dark room. The head, trunk, and legs were firmly secured to the seat with the bands and a four-point safety belt in a natural position (Fig. 1). An axis under the tilt-table was expanded or contracted via a servo motor, enabling the tilt table to be tilted in the roll plane around a rotation center located 18 cm underneath the bottom of the seat. The tilting velocity and initial acceleration were respectively 0.44°/s and 0.09°/s^2^, which is below the rotational acceleration threshold [33]. Therefore, in the present study, the contribution of the semi-circular canal to the estimation of visual vertical was negligible.

**Fig. 1.**
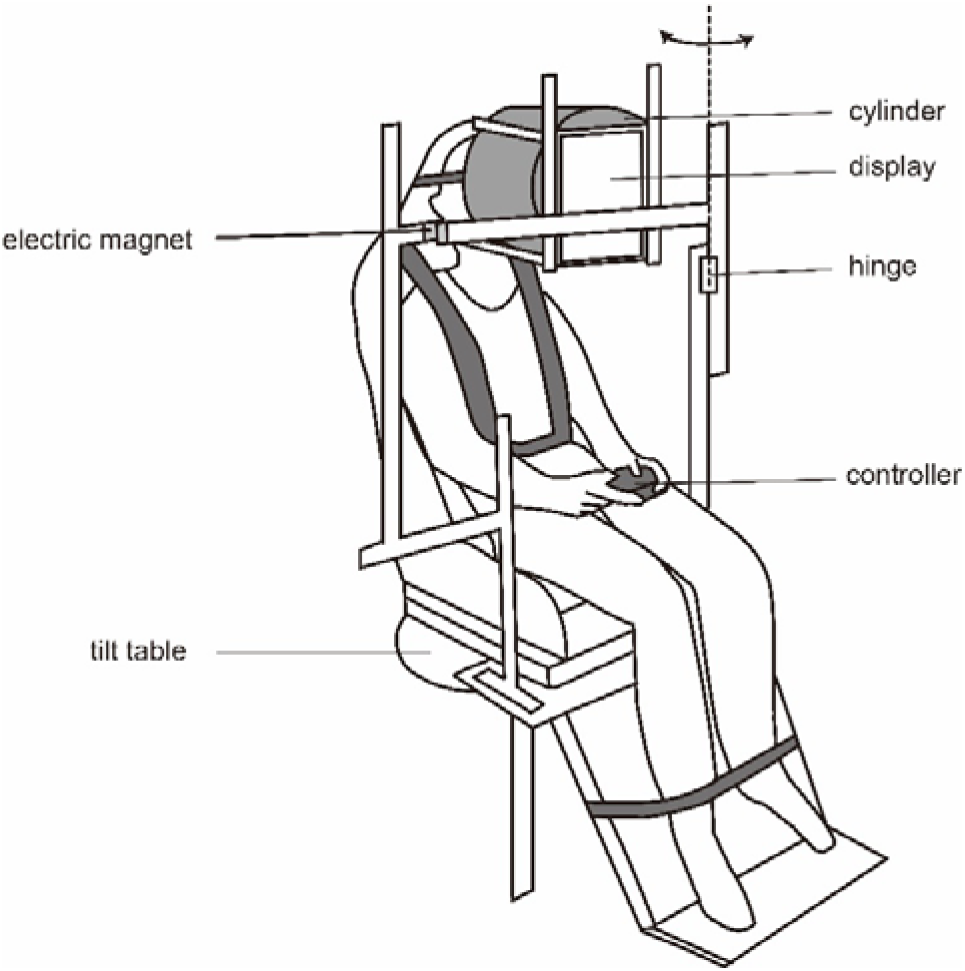
Schema of the experimental setup. The display portion could be rotated around the hinge joint only when the electric magnet was not electrified.

A display (LTN097QL01, SAMSUNG, Korea; 19.6 cm × 14.7 cm) was placed 35 cm in front of the participant’s face. To prevent any spatial cues such as the edge of the display, a black-colored cylinder (26 cm in diameter) with one end covered by a plate with a hole (10 cm in diameter) in the center was placed between the face and the display. During the SVV adjustment, a white line (length, 4 cm; width, 0.1 cm) which could be rotated via a digital controller (BSGP1204, iBUFFALO, Japan) was presented in the center of the display. An anti-aliasing mode was applied for the projection to avoid any orientational cues derived from the pixel alignment. The display was mounted on the tilt table via metal frames (Green Frame, SUS, Japan), maintaining the display position identical relative to the participants regardless of body tilt angle. At the center of a vertical frame positioned on the left side of the tilt table was the hinge structure, enabling it to rotate the display portion in the horizontal plane independently from the tilting chair. Before participants performed the task (see *Task during prolonged tilt* in detail), an experimenter rotated the display portion to the left side of the participants, preventing them from hitting their arm against the display or frames. An electric magnet was set between the display portion and the horizontal frame positioned at the right side of the tilt table. The display portion and frame were firmly fixed via the electrification of the electric magnet, enabling it to be set in front of the face. Prior to each experiment, the angle of the tilt-table and the upper side of the display relative to the floor was calibrated at 0° using a digital inclinometer, enabling us to accurately define the gravitational vertical in the SVV adjustment.

To temporarily restrict vision, the participants wore a mechanical shutter goggle controlled via microcomputer (Arduino UNO, Arduino SRL, United States) during the experiment. They also wore earphones via which white noise was provided to avoid auditory cues from the environment.

#### Experimental Procedure

This experiment consisted of two sessions, *during-tilt* and *post-tilt* sessions. In the former, we evaluated how the SVV was influenced by arm movements at the tilt position. In the latter, we confirmed whether the effects of arm movements during prolonged body tilt influenced the SVV after returning to the around upright position. The order of each session was randomized for each participant.

Figure 2A shows the experimental procedure in the *during-tilt* session. After the experimenter announced the start of the trial, the shutter closed and the tilt table was tilted leftward. One second later, after the tilt table came to the left-side-down (LSD) 16° position, the shutter opened again, and a white line was presented on the display. The participants were asked to adjust the line to the gravitational vertical via the controller (SVV adjustment). The initial angle of the line was set at ±45°, ±60°, or 90° relative to the body longitudinal axis in a pseudorandomized order. The participants repeated 5 trials within 40 seconds. Then the shutter closed and the display portion was moved leftward by the experimenter. The participants were asked to execute one of three tasks (see the *Task during prolonged tilt* section) at the tilt position. After the display portion was returned to the initial position (i.e. in front of the participant’s face), the shutter opened and the participants were asked to perform the SVV adjustments again for five trials. After the SVV adjustments, they were tilted back to upright. Approximately two minutes of rest was given between trials. Each participant performed the SVV adjustments for 30 trials [3 task conditions × 2 phase (before and after task) × 5 trials].

**Fig. 2.**
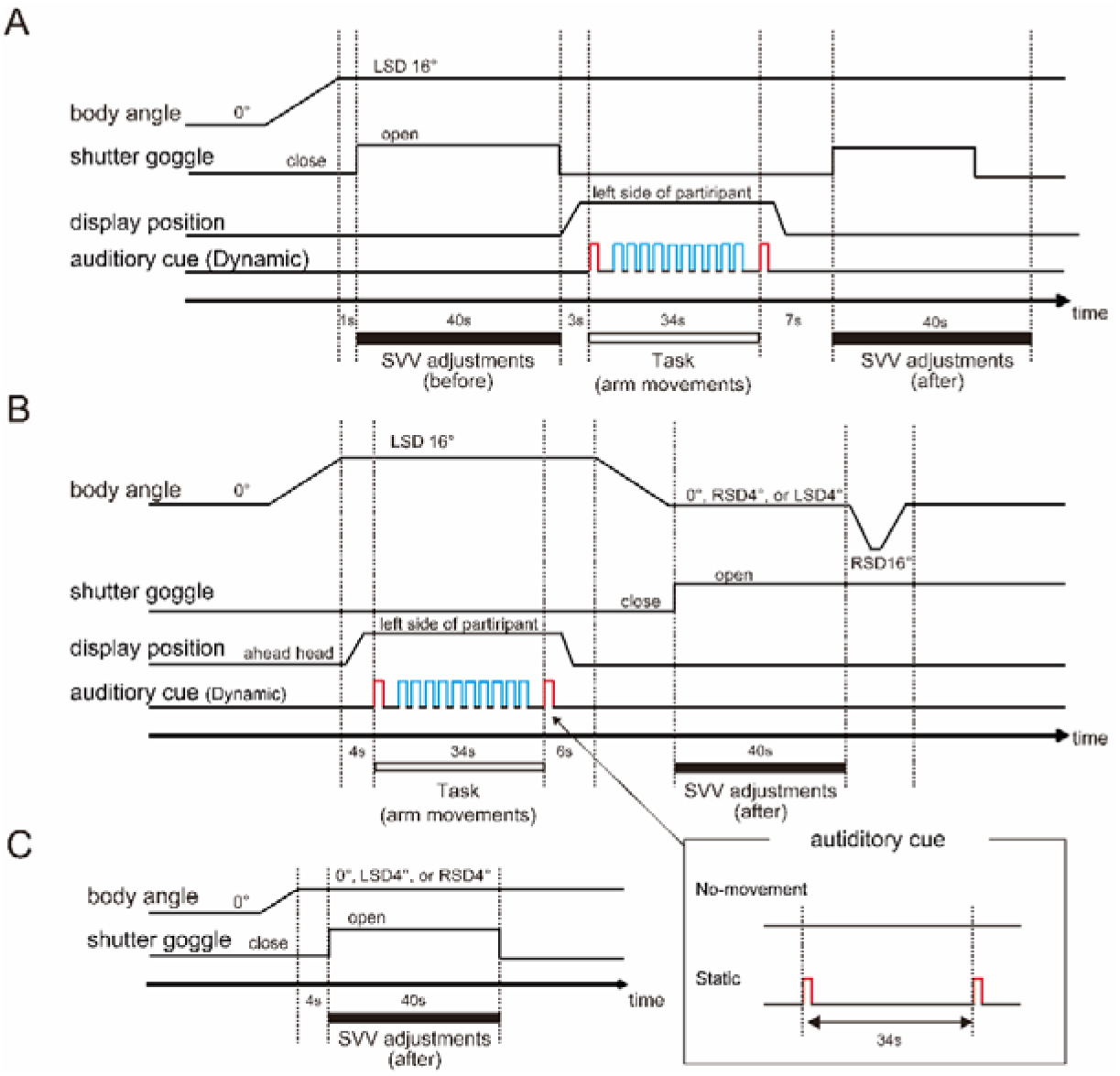
Schema of the experimental procedure in the *during-tilt* (A) and the *post-tilt* sessions (B). In both sessions, participants were asked to perform one of three tasks (No-movement, Static, or Dynamic task) during prolonged tilt in response to preparation (denoted as *red-colored*) or action sounds (denoted as *blue-colored*). As an example, the auditory cue for the Dynamic task is shown in the main figures. (C) Illustration of the Control condition in the *post-tilt* session. The participants performed the SVV adjustments immediately after arriving at one of the final tilt positions from the starting (upright) position.

Figure 2B shows the experimental procedure in the *post-tilt* session. After the participants were tilted to LSD 16° position from upright with the shutter closed, the display portion was moved leftward by the experimenter and the participants were asked to perform one of the three tasks during prolonged tilt (34s). Then, the display portion was then moved back to the original position and the tilt table was tilted to one of the following positions: upright, right-side-down (RSD) 4°, or LSD 4°. The shutter opened and the participants were asked to repeat the SVV adjustments for five trials. Then, the tilt table was tilted back to upright via RSD 16° to avoid providing feedback about body angle to the participants. To prevent the participants from performing the SVV adjustments based on prior knowledge about the experimental protocol, they were not informed about their final tilt position. Each participant performed the SVV adjustments for 45 trials [3 task conditions × 3 final tilt positions × 5 trials]. The order of the task conditions was pseudorandomized for each participant.

After all trials were accomplished, participants performed SVV adjustments for 5 trials at each final tilt position (0° and ±4°) immediately after being tilted from an upright position (not via the LSD position), referred to as the Control condition (Fig. 2C). Note that the effect of prolonged tilt (including the initial tilt) and of arm movements at the LSD 16° position would not be reflected on the angle of SVV in the Control condition.

#### Task during prolonged tilt

The participants performed one of three tasks during prolonged tilt as follows: No-movement, Static, or Dynamic tasks. For the No-movement task, both preparation and action sounds were not presented, and the participants were asked to just maintain the tilt posture. For the Static task, a preparation sound was first presented via the earphones, prompting the participants to switch the controller to their left hand and to point to the front of the face using their right index finger with the right arm extended. The participants were instructed to maintain this posture until another preparation sound was presented. For the Dynamic task, a preparation sound was first presented and participants were asked to set the pointing posture as with the Static condition. Three seconds after the preparation sound, an action sound was presented every 3 seconds for a total of 10 times. The participants were asked to move their arm upward and then down parallel to their body’s longitudinal axis once per one action sound with their arm extended. The length of arm movement was set from the height of the eye to the navel. Then, another preparation sound was presented, prompting them to hold the controller again.

Prior to the beginning of the experiments, participants sufficiently practiced each task. The duration time of each action condition (i.e. duration of prolonged tilt) was identical across all conditions (34 seconds). The order of the tasks was pseudorandomized across participants.

#### Data Analysis

We calculated SVV angle as a deviation between the subjective vertical and the actual gravitational vertical for each trial. The median of 5 trials was applied as the representative value in each task condition for each participant. To determine the extent of SVV shifts induced by prolonged tilt and arm movements, we calculated the *ΔSVV* values by subtracting the SVV angle before the task from the SVV angle after it for the *during-tilt* session, and by subtracting the SVV angle in the Control condition from the SVV angle in each task condition (No-movement, Static, Dynamic) for the *post-tilt* session. Results of the Shapiro-Wisk tests showed that SVV angles and *ΔSVV* values were normally distributed across participants in all conditions for the *during-tilt* session (all *p* > 0.05), but not in some conditions for the *post-tilt* session (*p* < 0.05). Therefore, we applied parametric statistical analyses to the dataset in the *during-tilt* session and non-parametric analyses to the dataset in the *post-tilt* sessions. Specifically, we compared the *ΔSVV in* each task condition by conducting one-way analysis of variance [ANOVA; 3 task conditions (No-movement, Static, Dynamic)] with repeated measures for the *during-tilt* session and Friedman tests [3 task conditions (No-movement, Static, Dynamic)] for the dataset in each tilt position (0, LSD or RSD 4°) for the *post-tilt* session.

For the ANOVA, the degree of freedom was corrected using Greenhouse-Geisser correction coefficient epsilon, and the *p*-value was recalculated if sphericity was found to be violated by Mauchly’s sphericity test. The significance level for all comparisons was set to 0.05. Bonferroni corrections were used for post-hoc multiple comparisons. All statistical analysis were conducted using R software version 3.5.3 (R Core Development Team, Austria).

### Experiment 2

#### Participants

Twelve right-handed healthy volunteers (8 males and 4 females, aged 22–26 years) participated in this experiment after giving written informed consents. As well as Experiment 1, this experiment was approved by the Ethics Committee of the Graduate School of Human and Environmental Studies, Kyoto University, and was conducted in accordance with the Declaration of Helsinki (2013).

#### Apparatus

As in Experiment 1, the participants sat on the tilting chair, and the head, trunk, and legs were firmly fastened to the seat. In this experiment, the velocities of the tilt table were different from in Experiment 1. The velocity of the tilt-chair during the subjective postural vertical (SPV) task (see *Procedure* paragraph) was set at 1.0°/s (initial acceleration: 0.52°/s^2^) to avoid the stimulation of semi-circular canals [33]. The velocity of the chair from upright to LSD 16° (before SPV task) and from RSD 16° to upright (after SPV task) was relatively fast (8.0°/s). However, based on a previous study [34] reporting that the amplitude of post-rotatory nystagmus was small even after roll body tilt with a speed of 10°/s, the contribution of the semicircular canal to SPV estimations was negligible.

The participants held a custom-made controller with a press-button to indicate the position of the perceived body vertical. The roll-tilt angle of the chair was monitored via an accelerometer module (KXM52-1050, Kionix, U.S.A.) mounted under the center of the tilt table. The signals from the accelerometer and from the controller were recorded via a data acquisition system (Power Lab 16sp, AD Instruments, Australia). The sampling frequency was set at 100Hz.

During the experiment, the participants wore an eye-mask and were provided white noise via earphones so as not to provide visual or auditory cues from the environment. To prevent the participants from being fatigued, a rest for about ten minutes was inserted per ten SPV trials.

#### Experimental procedure

The blindfolded participants were first tilted to LSD 16°. In this position, they were presented with one of four task conditions (No-movement, Static, Dynamic, Control). Under the former three conditions, as with Experiment 1, they were asked to perform each task based on the preparation and action sounds (see *task during prolonged tilt*), and then they were tilted to RSD 16°. While tilted from LSD 16° to RSD16°, they were asked to press the bottom of the controller when they felt body was upright (SPV task). In the Control condition, they were tilted to RSD 16° immediately after arriving at LSD 16° (i.e. without prolonged tilt) and performed the SPV task. After each SPV task, they were tilted back toward upright. As with Experiment 1, the duration time of each task was 34 seconds.

All the participants performed 7 trials of the SPV task for each of the 4 task conditions, i.e., a total of 28 trials. The order of presentation of the task conditions was pseudorandomized for each participant.

#### Data Analysis

We calculated SPV angle as the deviation between the subjective vertical and actual gravitational vertical for each trial. The median was used as the representative value of SPV angles in each task condition for each participant. Similar to SVV, we quantified the extent of SPV shifts (*ΔSPV*) induced by prolonged tilt and arm movements by subtracting SPV angle in the Control condition from those in each task condition (No-movement, Static, and Dynamic). Since the SPV angles in each task condition were normally distributed across participants (Shapiro-Wisk tests, *p* > 0.05), we conducted one-way ANOVA with repeated measures [3 task conditions (No-movement, Static, Dynamic)] to compare *ΔSPV* values between task conditions. If sphericity was found to be violated by Mauchly’s sphericity test, the degree of freedom was corrected using Greenhouse-Geisser correction coefficient epsilon, and the p-value was recalculated.

## Results

### Experiment 1

#### Prolonged tilt effect

We first checked whether the prolonged tilt induced the SVV shifts in the present experimental set-up. Figure 3 shows the angular changes of SVV in the No-movement condition in each session. Positive and negative values of SVV correspond to rightward and leftward deviation, respectively. For the *during-tilt* session, a paired t-test revealed SVV angle after No-movement task (1.3 ± 1.2°) significantly shifted leftward, compared to the result before its task (−0.4 ± 1.3°; *t*_11_ = 2.58; *p* < 0.05, cohen’s *d* = 0.67). For the *post-tilt* session, Wilcoxon signed ranked tests showed that SVV angle in the No-movement task significantly shifted leftward, compared to the Control condition for LSD 4° (*p* < 0.001, effect size *r* = 0.86) and 0° positions (*p* < 0.05, effect size *r* = 0.78), but not for RSD 4° position (*p* = 0.35, effect size *r* = 0.28).

**Fig. 3.**
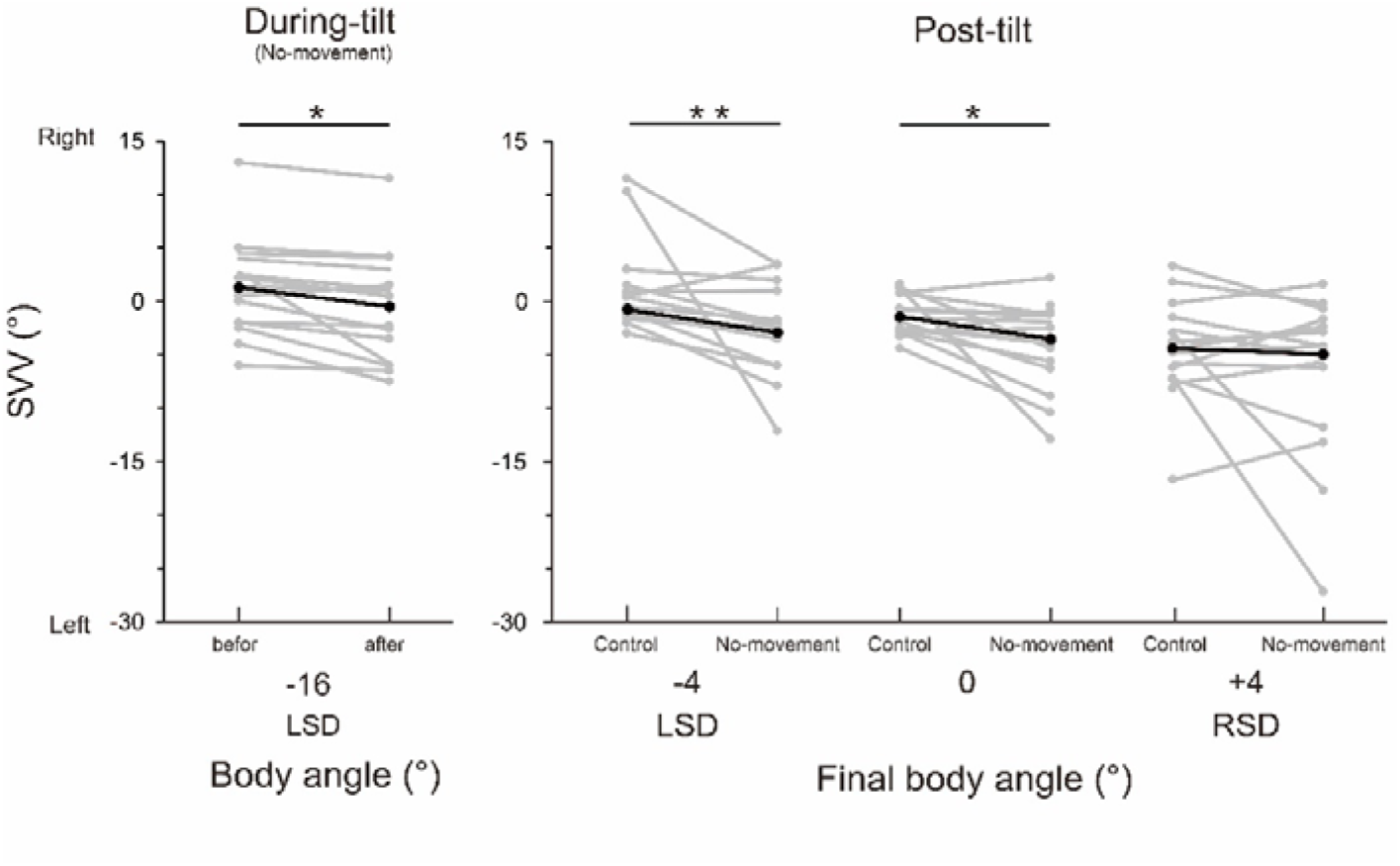
The alteration of SVV angles in the No-movement condition. Grey bars denote the individual median data and black bars denote the group-mean (*during-tilt* session) or −median data (*post-tilt* session). *: *p* < 0.05, **: *p* < 0.01

#### During-tilt session

Figure 4A shows group-mean *ΔSVV* value in each task condition. The one-way ANOVA results revealed a significant main effect of task condition (*F*_*2*, 28_ = 4.77, *p* < 0.05, effect size *η^2^* = 0.14). Results of the post-hoc Bonferonni tests showed that *ΔSVV* in the No-movement (−1.6 ± 0.6°) and Static conditions (−1.2 ± 0.4°) were smaller (i.e., shifted leftward), compared to the Dynamic condition (0.2 ± 0.4°; vs No-movement, *p* < 0.05, effect size *r* = 0.60; vs Static, *p* < 0.05, effect size *r* = 0.54). Significant differences were not noted between the Static and No-movement conditions (*p* = 0.46, effect size *r* = 0.19). These results indicate that the SVV shifts occurred during prolonged tilt were attenuated when performing dynamic arm movements.

**Fig. 4.**
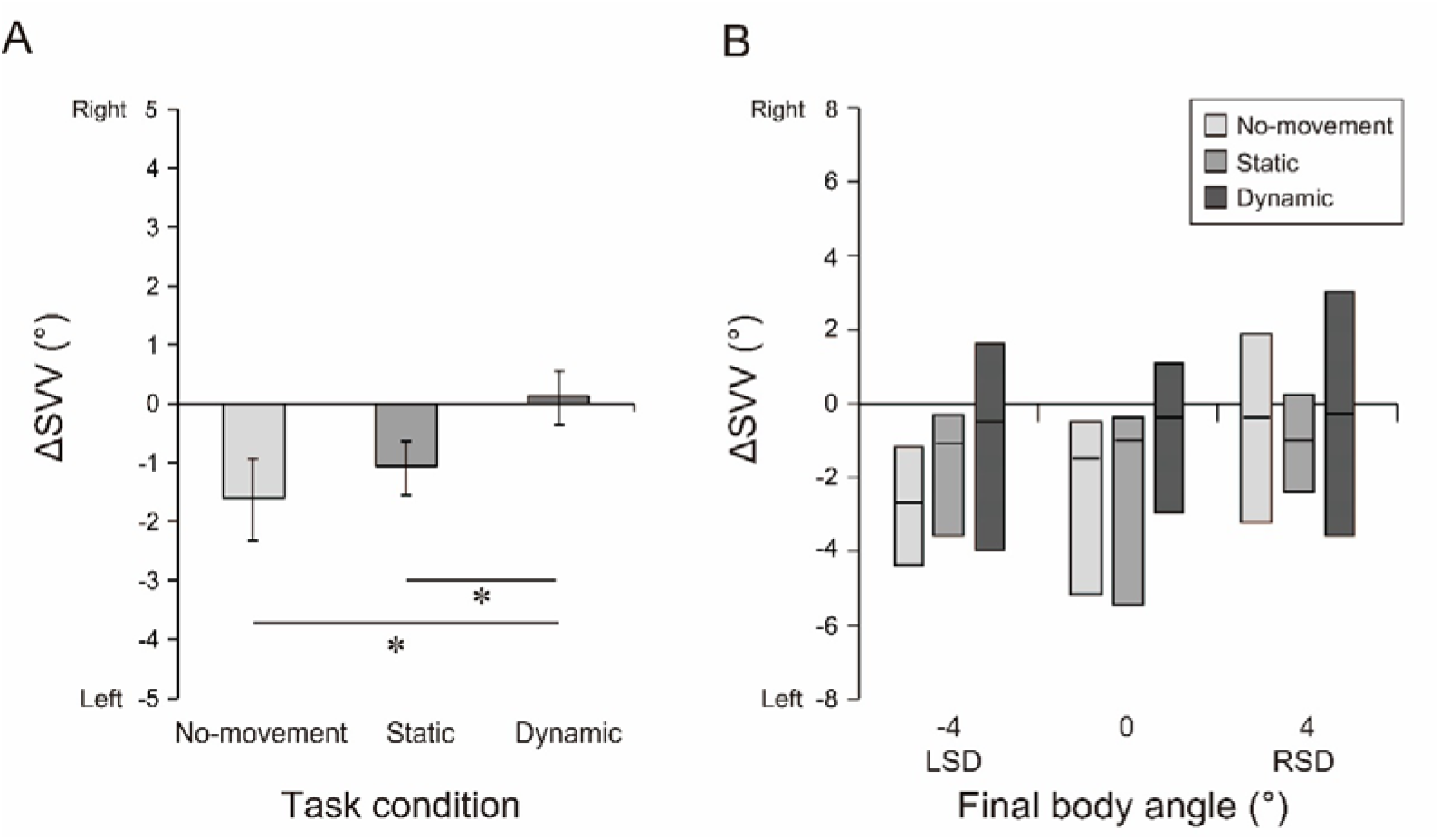
**(A)** The group-mean *ΔSVV* values in the *during-tilt* session. Error bars denote standard errors across participants. **(B)** The group-median *ΔSVV* values in the *post-tilt* session. The horizontal line within each box, and the lower and upper ends of each box represent median, and 1st and 3rd quartiles, respectively. *: *p* < 0.05

#### Post-tilt session

Figure 4B shows group-median *ΔSVV* in each action condition at each final tilt position. Results of the Friedman tests revealed a significant main effect of task condition for LSD 4° position (*χ^2^* = 8.40, *p* < 0.05), but not for 0° (*χ^2^* = 2.53, *p* = 0.28) and RSD 4° (*χ^2^* = 0.85, *p* = 0.65). For the LSD 4° position, however, results of the post hoc tests showed no significant differences in *ΔSVV* among different task conditions (No-movement vs Dynamic, *p* = 0.13, effect size *r* = 0.58; No-movement vs Static, *p* = 0.15, effect size *r* = 0.52; Static vs Dynamic, *p* = 0.44, effect size *r* = 0.38). These results indicate that the SVV shifts occurred after prolonged tilt were not clearly influenced by either static or dynamic arm movements during prolonged tilt.

### Experiment 2

Figure 5A shows the mean SPV angles in the Control and No-movement conditions. The SPV angle significantly shifted leftward in the No-movement condition (−5.0 ± 0.9°), compared to the Control condition (−0.8 ± 1.2°; *t*_11_ = 5.77, *p* < 0.001, cohen’s *d* = 1.67). This result indicates significant SPV shifts induced by prolonged tilt.

**Fig. 5.**
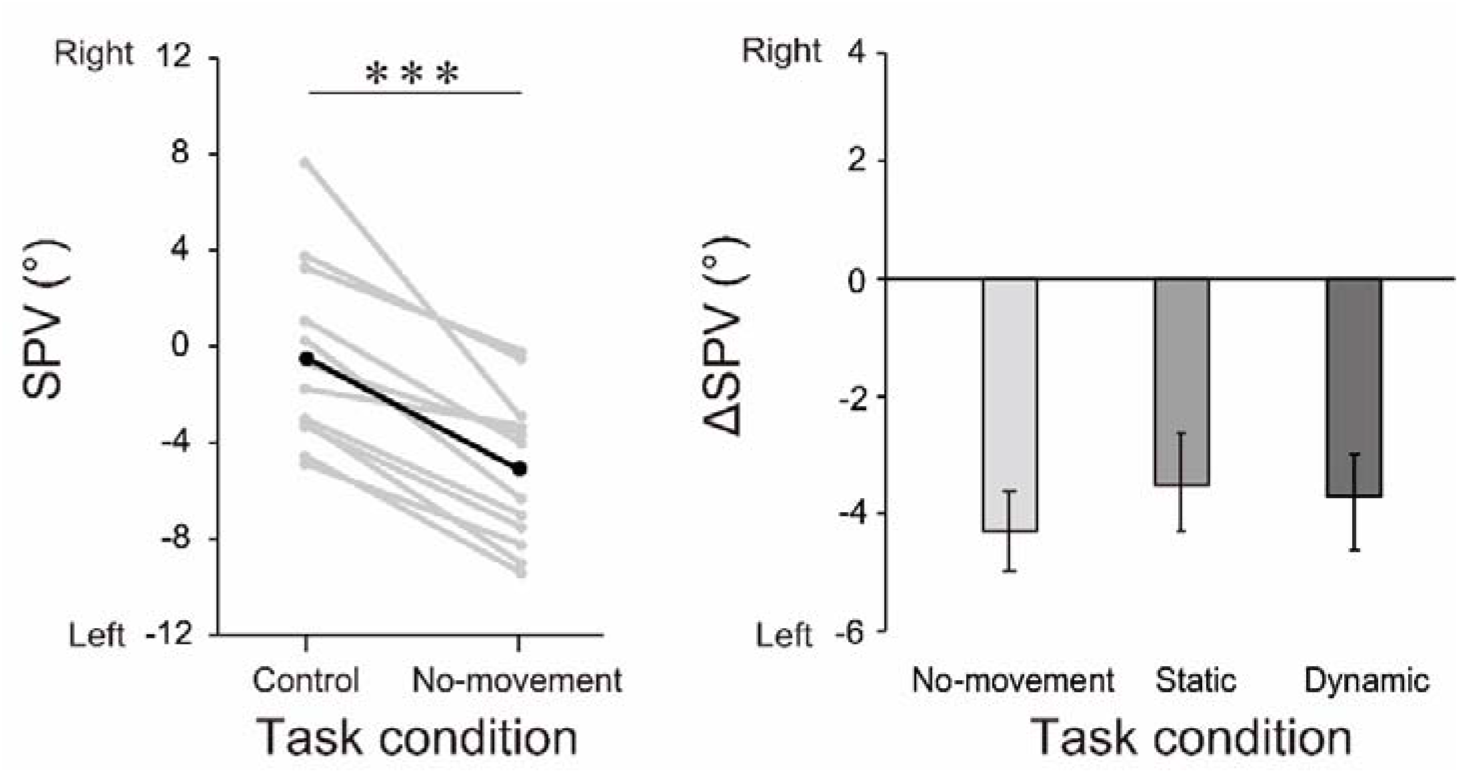
**(A)** SPV angles in the Control and No-movement conditions. Gray and black lines represent individual median and group-mean values, respectively. **(B)** The group-mean *ΔSPV* value in each task condition. Error bars represent standard errors. ***: *p* < 0.001

Figure 5B shows the group-mean *ΔSPV* values in each task condition. The mean (±SE) *ΔSPV* were −4.3 ± 0.7° for the No-movement condition, −3.5 ± 0.8° for the Static condition, and −3.9 ± 0.8° for the Dynamic condition, respectively. The one-way ANOVA results revealed a non-significant main effect of the task condition (*F*_*2*, 22_ = 2.54, *p* = 0.10) with small effect size (*η*^2^ = 0.01). This result indicates that neither static nor dynamic arm movements during prolonged tilt influenced the prolonged tilt-induced SPV shifts.

## Discussion

The present study investigated how static or dynamic arm movements altered the effect of prolonged tilt on performances in the SVV and SPV tasks. In Experiment 1, we found that the performance of dynamic arm movements effectively attenuated the SVV shifts occurred during prolonged tilt (*during-tilt* session), but not those after prolonged tilt (*post-tilt* session). In experiment 2, SPV angles were not clearly affected by either static or dynamic arm movements.

We confirmed the significant SVV shifts toward the side of body tilt in the *during-tilt* session (Fig. 3, *left panel*) consistently with results reported in the literature [17–22], but these SVV shifts decreased when the participants performed dynamic arm movements during prolonged tilt (Fig. 4A). Given that the SVV shifts induced by prolonged tilt would be mainly due to sensory adaptations in the vestibular and body somatosensory systems [11,28], these results suggest that the CNS would utilize not only vestibular or static body somatosensory signals, but also the additional cues generated by dynamic body movements for estimating the visual vertical.

Conversely, we observed no clear effects of static arm movements on SVV in the *during-tilt* session, even though the gravitational force on the arm—dependent on the body’s tilt angle—is generated by both static and dynamic arm movements. The lack of an effect of static arm movements may imply that the dynamic property of arm movements is important for estimating the visual vertical. The gravitational force on the arm during arm movements can be perceived as a sense of heaviness based on the afferent information about muscle tension from the Golgi tendon organ (GTO) located at the muscle-tendon junction [35,36]. Psychophysical studies have shown that estimation of the heaviness of an object with concurrent dynamic movements, such as lifting or wielding, is more accurate than estimation with static holding [37,38]. In addition, a physiological study has demonstrated that when constant tension is persistently applied to a muscle, the firing rate and the sensitivity of the GTO to the force gradually deteriorate [40]. These findings lead us to speculate that information about the gravitational force on the arm might be conveyed to the CNS more accurately while performing dynamic rather than static arm movements, leading to attenuation of the prolonged-induced SVV shifts.

In addition to the above-mentioned assumption, the possibility that the neck muscle activities accompanying arm movements influenced the SVV must be also considered. Although the head and trunk were firmly secured to the tilting chair with a seatbelt and bands, we cannot disregard the fact that dynamic arm movements may have resulted in increased or decreased activities at the neck muscles. Due to the involvement of neck proprioception in visual vertical estimation [40,41], neck muscle activities during dynamic arm movements may have partially been responsible for the attenuation of the prolonged tilt-induced SVV shifts.

In the *post-tilt* session, significant SVV shifts after prolonged tilt were observed at the final tilt position of LSD 4° and 0° (Fig. 3, *right panel*). However, in contrast to the *during-tilt* session, the dynamic arm movements did not effectively attenuate these SVV shifts (Fig. 4B). A potential explanation for this difference between the sessions may be attributable to the duration after the arm movements. In the *during-tilt* session, the participants performed SVV adjustments immediately after the arm movement task. On the other hand, in the *post-tilt* session, they performed SVV adjuestments after slowly tilting back toward each final tilt position, thus the interval between dynamic arm movements and SVV adjustments was relatively long (at least 20s). The contribution of the additional cues derived from dynamic arm movements to visual vertical estimates likely diminished over time after the task, thereby resulting in no clear effects of dynamic arm movements on SVV in the *post-tilt* session.

The prolonged tilt-induced SPV shifts were not effectively attenuated by either static or dynamic arm movements (Fig. 5). Due to the relatively long time between the estimation of postural vertical and the arm movement task, as well as the *post-tilt* session, the lack of clear effect of arm movements on the SPV may also be attributed to the temporal decay of the arm movement effects.

This study consists of a limitation—the sample size is relatively small. In particular, at LSD 4° position in the *post-tilt* session, no significant differences were noted between the No-movement condition, and the Static or Dynamic conditions, despite the large effect size (*r* > 0.50), which is likely a result of the small sample size. Future studies must include a larger sample size to conclude the effect of arm movements on the SVV shifts after prolonged tilt.

## Conclusion

The present study shows that dynamic arm movements can attenuate the SVV shifts that occur during prolonged tilt. This finding suggests that the supplementary information generated by dynamic body movements plays an important role in the perception of visual vertical as well as vestibular and body somatosensory signals. To provide an improved understanding of the relationship between the effects of body movements on the perception of the gravitational space, we need to further examine how performance in the estimation of the gravitational direction would be influenced by the manipulation temporal (e.g., arm movement velocity, interval between arm movements and perceptual task) and spatial properties (e.g., direction of arm movements or body tilt).

